# Understanding 6th-Century Barbarian Social Organization and Migration through Paleogenomics

**DOI:** 10.1101/268250

**Authors:** Carlos Eduardo G. Amorim, Stefania Vai, Cosimo Posth, Alessandra Modi, István Koncz, Susanne Hakenbeck, Maria Cristina La Rocca, Balazs Mende, Dean Bobo, Walter Pohl, Luisella Pejrani Baricco, Elena Bedini, Paolo Francalacci, Caterina Giostra, Tivadar Vida, Daniel Winger, Uta von Freeden, Silvia Ghirotto, Martina Lari, Guido Barbujani, Johannes Krause, David Caramelli, Patrick J. Geary, Krishna R. Veeramah

## Abstract

Despite centuries of research, much about the barbarian migrations that took place between the fourth and sixth centuries in Europe remains hotly debated. To better understand this key era that marks the dawn of modern European societies, we obtained ancient genomic DNA from 63 samples from two cemeteries (from Hungary and Northern Italy) that have been previously associated with the *Longobards*, a barbarian people that ruled large parts of Italy for over 200 years after invading from Pannonia in 568 CE. Our dense cemetery-based sampling revealed that each cemetery was primarily organized around one large pedigree, suggesting that biological relationships played an important role in these early Medieval societies. Moreover, we identified genetic structure in each cemetery involving at least two groups with different ancestry that were very distinct in terms of their funerary customs. Finally, our data was consistent with the proposed long-distance migration from Pannonia to Northern Italy.

## INTRODUCTION

Western Europe underwent a major socio-cultural and economic transformation from Late Antiquity to the Early Middle Ages. This period is often characterized by two major events: the collapse of the western Roman Empire, and its invasion by various western and eastern barbarian peoples such as the Goths, Franks, Anglo-Saxons, and Vandals, as well as by nomadic Huns; as such it has come to be known as the Migration Period, in German the *Völkerwanderung,* and in French *Les invasions barbares*. However, written accounts of these events are laconic, stereotypical, and largely written decades or even centuries later^1,2^. Because barbarian populations of the Migration Period left no written record, the only direct evidence of their societies comes from their archaeological remains, chiefly grave goods, that have been used to make inferences about group identities, social structures, and migration patterns^3–5^. Interpretation of the archaeological record is also fraught with difficulties. Grave goods represent a limited and highly curated portion of material culture, and with little other archaeological data available fundamental questions about barbarian social organization remain unanswered^6^. As such, almost all aspects describing the people and events of this period are still a matter of vigorous debate^7,8^. Were specific barbarian peoples described in texts culturally and ethnically homogeneous populations with well-defined social and political institutions, or were they ad-hoc and opportunistic confederations of diverse, loosely connected groups? Did migrations involve tens or hundreds of thousands of individuals as described by the Romans or simply small military units of male warriors? How rapidly did they mix culturally and genetically with other barbarian populations and with the resident populations within the Empire in the provinces that they came to dominate?

One group with a relatively voluminous historical description are the Longobards (also known as Lombards, Longobardi or “Long-beards”^9–11^). According to written sources, the Longobards were barbarians first identified as living east of the lower Elbe River in the first century CE. After disappearing from written sources for three centuries, Longobardi are reported around 500 CE north of the Danube, from where they then expanded into the Roman province of Pannonia (what is now Western Hungary and Lower Austria). In 568 CE the Longobard King Alboin led an ethnically mixed population into Italy, where they established a kingdom covering much of the country until 774 CE (Fig 1A). Numerous archaeological sites in Pannonia and Italy contain broadly similar grave goods and burial customs, a pattern consistent with the historical account of a Longobard migration into Italy (on history of the Longobard tradition, see Supplementary Text S1). One of the few contemporary texts describing the movement of the Longobards is by the Roman bishop Marius of Avenches who states “*Alboin king of the Longobards, with his army, leaving and burning Pannonia, his country, along with their women and all of his people occupied Italy in fara*”. Etymologically the word *fara* is rooted in ‘to travel”, but its precise meaning is ambiguous. While some have interpreted it to represent cognatic, kin-based clans, others suggest that it may simply refer to military units of mixed background^12^.

**Fig 1.**
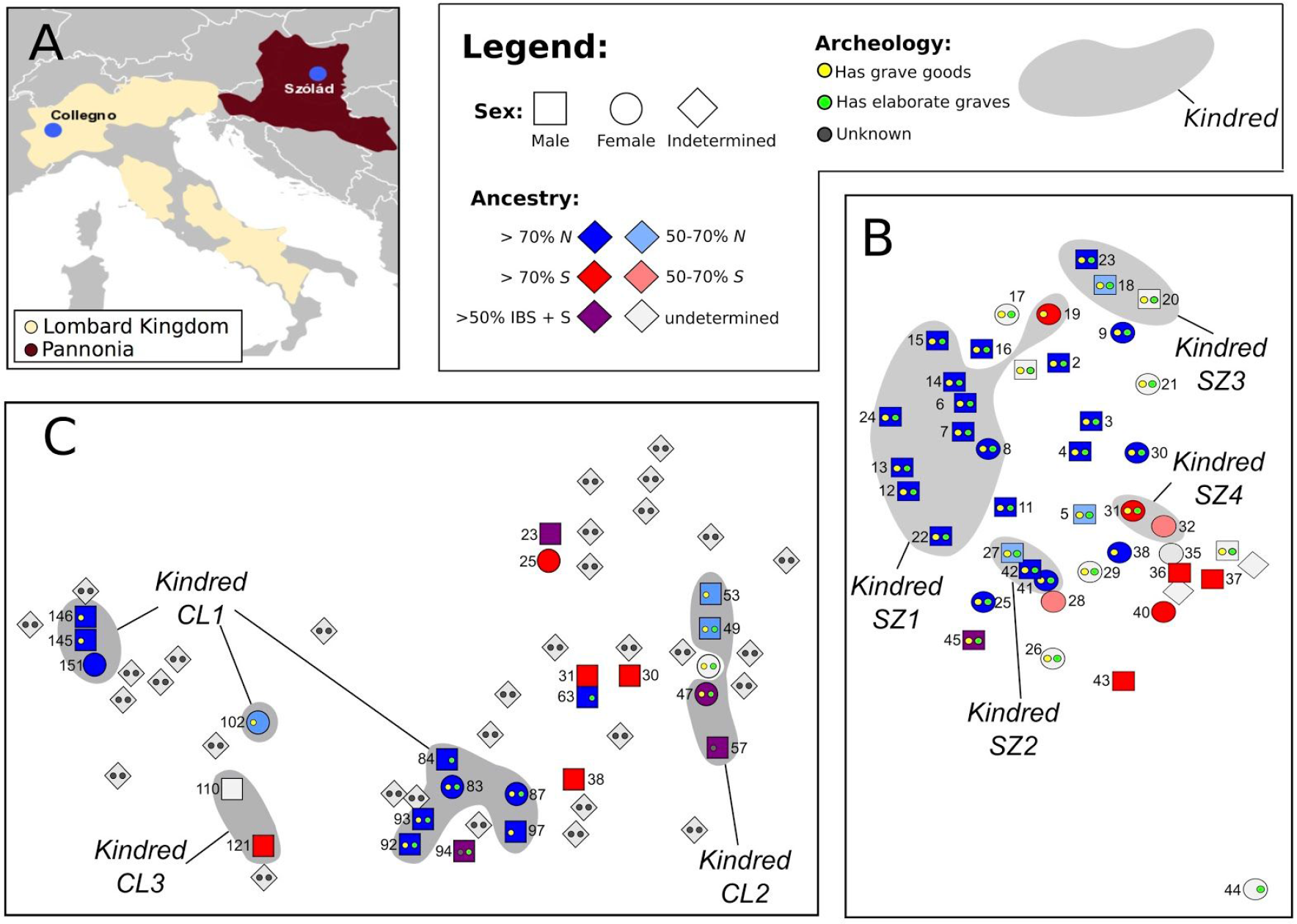
Archaeological and genetic characterization of Szólád and Collegno. (A) Map of Europe showing the location (blue dots) of the two cemeteries and regional context is included (the Roman province of Pannonia in burgundy and the Longobard Kingdom in beige). (B) and (C) show the spatial distribution of graves in Szólád and Collegno (first period burials only) with indication of sex (different shapes), genetic ancestry (different colors) and summary of archeology (yellow dots for presence/absence of grave furnishings and green dots for the presence of wooden elements in grave structure). Kindreds (in the biological sense) are indicated by gray shading in panels (B) and (C). N = FIN+GBR+CEU, S = TSI.

In this study we contribute new evidence to the debate surrounding Migration Period barbarians by generating paleogenomic data for 63 individuals from two key sixth to seventh century cemeteries associated with the Longobards, Szólád in western Hungary and Collegno in Northern Italy (Fig 1A). We note that we are not aiming to infer “Lombard ethnicity”, which is a subjective rather than objective identity. Our approach is unique in that we attempted to genomically characterize all of the interred individuals, rather than sampling specific individuals based on certain material culture markers. Combined with evidence of material culture, mortuary practices and isotope data, our approach provides an unparalleled image of the social organization of these historical communities, and begins to shed new light on possible movements within Europe during this period.

## RESULTS

### Two Longobard-associated cemeteries

We performed a deep genomic characterization of individuals buried in two 6th - 7th centuries CE cemeteries that have material culture associated with the Longobard era and the Longobards. Both are considered key sites with regard to the proposed migration from Pannonia to Italy. The first cemetery, Szólád, is located in present-day Hungary (Figure S2.1). There are 45 graves (Fig 1B), all of which are dated to the middle third of the sixth century based on a combination of stylistic elements of the grave goods and radiocarbon analysis^13^. Archaeological, stable isotope and mtDNA (HVS-1) analyses suggested that Szólád was occupied for only ~20/30 years by a mobile group of “Longobard era” settlers^13^. The female to male ratio (sexing being based primarily on genetic data, or in its absence, anatomy) is 0.65. Graves in this cemetery are organized such that a “core group” (N=18), mostly of male individuals, is surrounded by a “half-ring” of females (N=11) (Figure S2.2). Most of these individuals lie in elaborate graves with ledge walls and wooden chambers all in the same orientation, furnished with numerous artifacts such as beads, pottery, swords and shields. The remaining 16 Longobard-period graves are more diverse in relation to the sex of the individuals, as well as to the quality of grave construction and richness of artifacts. Archaeologists also recovered two bodies (AV1, AV2) that derive from a later occupation of the region by the Avars in the fill of the Longobard-period grave (SZ27)^14^, as well the skeletal remains from an individual dating to the Bronze Age 10m north of the cemetery (SZ1). See Supplementary Text S2 and Figures S2.3-11 for the archaeological context of Szólád.

The other cemetery, Collegno, is near Turin (Figure S3.1), northern Italy, and was in use from the late sixth through the eighth centuries, the earliest period of the Longobard kingdom in Italy^15^. We studied the 57 graves that date between 580 and 630 CE based on artefact typologies (Fig 1C) and represent the first of three major periods of occupation. The cemetery developed from the center outwards to the east and west. The types and range of grave goods in these 57 interments are comparable to those recovered at Szólád. However there is also evidence for a gradual cultural and religious evolution, with some practices disappearing in later decades. While there are no ledged graves, some are constructed via a wooden chamber structure, and there is the skeleton of a horse (devoid of head) in both cemeteries. See Supplementary Text S3 and Figures S3.2-11 for the archaeological context of Collegno.

Illumina sequencing of DNA extracts from petrous bone^16^ identified 38 and 21 samples from Szólád and Collegno, respectively, for which there was high endogenous content, high library complexity, and patterns of postmortem damage (PMD) characteristic of ancient DNA (Supplementary Text S4, Table S4.1). Another four DNA extracts (one from Szólád, three from Collegno) from teeth were also included in our dataset for samples lacking petrous bone DNA. Analysis of X-chromosome-mapped reads in males and mtDNA in both sexes revealed low levels of estimated contamination in almost all samples (mean ~1%), though one, CL31 had a value of 27% and 7% using X and mtDNA reads respectively. While we include this sample in certain individual-based analyses, its results should be treated with caution. Genomic libraries for the majority of samples (60 out 63) underwent partial UDG treatment^17^. Unique endogenous DNA content was sufficient for 10 male samples from Szólád, obtained from petrous bones, to undergo whole-genome sequencing (WGS), with a mean genome-wide coverage across samples of 11.3x. The remaining 53 samples underwent an in-solution capture protocol that targeted a set of previously described 1.2M single nucleotide polymorphisms (SNPs) (“1240k capture”)^18,19^. The average coverage at these SNPs (excluding the whole genomes) was ~1.5x and the mean number of genotyped SNPs per sample was ~522K (Supplementary Text S5). Unless noted, we considered 33 and 22 samples from Szólád and Collegno for downstream analysis, respectively (three and one samples from Szólád and Collegno had fewer than 30,000 usable SNPs, while samples SZ1, AV1, AV2 and CL36 were found based on archeology not to belong to the same occupation period as the other samples, as described above).

In addition, we assembled comparative 1240K capture SNP data for a number of different modern reference Eurasian samples, either using directly genotyped SNPs from WGS^20,21^ or by imputation^22,23^, as well as 211 ancient West Eurasians from 6,300 - 300 BC^19^ (Supplementary Text S6). We also included comparative genomic data from seven genomes from the UK associated with the Anglo-Saxons^24^ and two from Bavaria^25^ that are from similar time periods as the two cemeteries focused on here (i.e. the Migration Period).

### Genetic ancestry in both cemeteries involves two primary central/northern and southern groups

Principal Component Analysis (PCA) of our ancient samples against modern reference sets infers that our ancient samples possess genetic ancestry that overlaps overwhelmingly with modern Europeans (Fig 2A. Figures S7.1-3; Supplementary Text 7). However, rather than being clustered close to their respective modern countries of origin, samples from Szólád and Collegno can be placed along the major northern and southern axis of modern European genetic variation. This north/south axis of genetic variation is also observed when examining only our ancient samples based on a set of ~50K SNPs for which haploid calls with no missing data can be made amongst 20 and 10 unrelated (see below) individuals from Szólád and Collegno respectively, demonstrating that our results are not a bias introduced due to the reliance on modern reference populations or close kinship (Figures S7.8-10).

**Fig 2.**
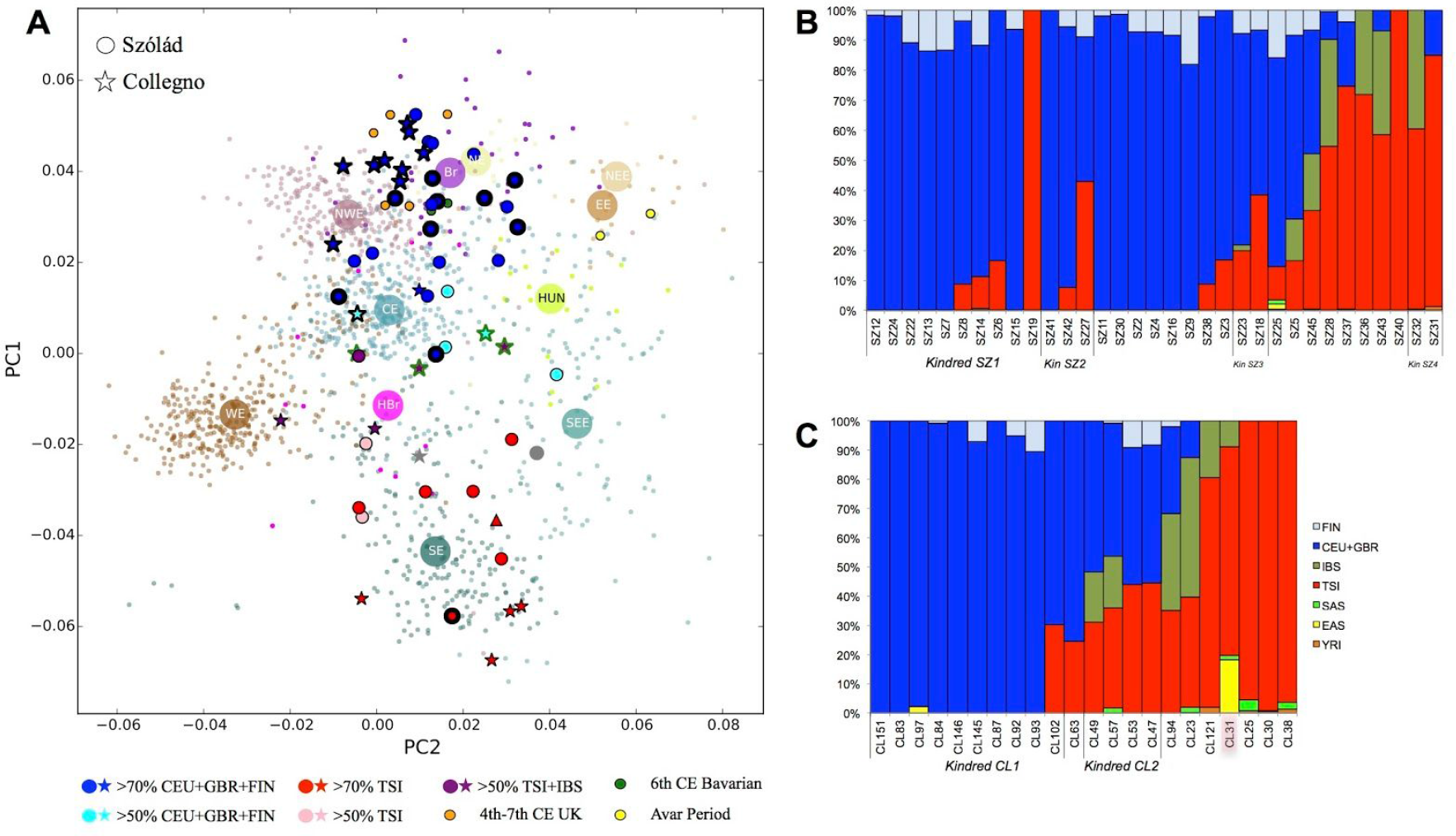
Genetic structure of Szólád and Collegno. (A) Procrustes Principal Component Analysis of modern and ancient European population (faded small dots are individuals, larger circle is median of individuals) along with samples from Szólád (filled circles), Collegno (filled stars), Bronze Age SZ1 (filled grey circle), second period CL36 (grey star), two Avar-period samples from Szólád (yellow circles), Anglo-Saxon period UK (orange circles) and 6th Century Bavaria (green circles). Szólád and Collegno samples are filled with colors based on estimated ancestry from ADMIXTURE. Blue circles with thick black edge = *Kindred_SZ1*, blue stars with thick black edge = *Kindred_CL1*, stars with thick green edge = *Kindred_CL2*. NWE = northwest Europe, NE = modern north Europe, NEE = modern northeast Europe, CE =central Europe, EE = eastern Europe, WE =western Europe, SE = southern Europe, SEE = southeast Europe, HUN = modern Hungarian, HBr = Hungarian Bronze Age, Br = central, northern and eastern Europe Bronze age. Model-based ancestry estimates from Admixture for Szólád (B) and Collegno (C) using 1000 Genomes Project Eurasian and YRI populations to supervise analysis. Note that high contamination was identified in CL31 and is shown with a triangle in the (A) and overlaid with a pink hue in the (C).

We next analyzed our ancient samples using supervised model-based clustering analysis implemented in ADMIXTURE^26^ (Supplementary Text 8), with Finns (FIN), British and Central Europeans (CEU+GBR, we found it difficult to consistently distinguish ancestry from these two populations), Tuscans from Italy (TSI), Iberians from Spain (IBS), South Asians (SAS), East Asians (EAS) and Yoruba (YRI) set as eight “parental” populations (Figures S8.1-6). CEU+GBR is the major genetic component in Szólád (mean of 64% across samples), with it being the major type in 25 out of the 33 individuals (Fig 2B). The second most prominent ancestry is TSI (average 25% across samples), and is the major component in the remaining nine individuals. Interestingly, despite being located in modern northern Italy, CEU+GBR ancestry is still the predominant type in Collegno, albeit at a smaller proportion (mean 57% across samples) and the major type in 15 of the 22 individuals, while TSI ancestry accounts for 33% across samples and is the major type in six individuals (Fig 2C). IBS ancestry is the major type in CL23 (48%) and is also prominent in CL94 (33%). By crudely assigning individuals into five colour-coded groups based on relative ancestry components, a clear correspondence can be observed between our ADMIXTURE and PCA analysis. Analysis of Y chromosomes in males generally reveals a highly concordant pattern to the autosomes, with haplogroups that are most predominant in modern central/northern Europeans along with a smaller set of southern European-associated haplogroups in both cemeteries (Supplementary Text 9, Figures S9.1-3, Table S9.2).

When conducting a Population Assignment Analysis (PAA) (Supplementary Text 10, Table S10.1) for each ancient sample against modern European reference populations, individuals with high TSI ancestry (>70%) from Collegno are primarily assigned to the Italian peninsula with high probability, while those from Szólád are more diverse, being assigned to both south and southeast Europe. Individuals with high CEU+GBR ancestry are assigned to countries from all over modern central and northern (including Scandinavia) and northwest Europe. We refer to this as central/northern ancestry as it is generally difficult to distinguish this with more precision given the resolution of our data. An analysis of rare variants in our nine medieval whole genomes from Szólád following the approach of Schiffels et al.^24^ (Supplementary Text S11) is consistent with the more common SNP-based analysis in terms of relative amounts of central/northern and southern ancestry (Figures S11.1-2). Five of the nine whole genomes are placed along extant European branches (four along the branch for modern Danes and one for TSI), and the remaining samples are placed on internal branches that connect modern northern and southern European populations (Figures S11.3-4).

We also examined our ancient samples within the context of the prehistoric groups that were the major contributors to modern European genetic variation: Paleolithic hunter-gatherers (WHG), Neolithic farmers (EEF) and Bronze Age Steppe herders (SA). Both PCA (Figures S7.4-7) and supervised ADMIXTURE analysis (Figures S8.16-18) essentially reiterate the same structure amongst our ancient samples, with greatest EEF ancestry in those individuals demonstrating similarities to modern southern Europeans and greater WHG+SA ancestry in those that resembled modern northern Europeans (with WHG being predominant in Northwest Europe and SA in Northeast Europe). Most individuals from Szólád (and Collegno) are genetically most similar not only to modern central/northern Europeans and 4-7th century UK and Bavarian genomes, but also to Bronze Age northern Europeans, rather than either modern or Bronze Age Hungarians (considering the genomes examined in Mathieson et al.^19^ as well as SZ1, the Bronze Age individual sampled at the Szólád site itself). Even the two later Avar-era genomes found in grave 27 in Szólád are genetically differentiated from the Lombard-era burials (they are genetically most similar to modern eastern Europeans), suggesting rapid population turnover in the region and the site itself.

### Both cemeteries are organized around close biological relatedness

We took advantage of genome-wide SNP information to infer pairwise biological relatedness within ancient cemeteries at an unprecedented level compared to previous studies^27,28^. We hereafter refer to groups of biologically related individuals as “kindred” as a shorthand, though we recognize that “kinship” in the traditional archaeological and anthropological sense encompasses a much broader range of social relationships^29^. We utilized lcMLkin^30^, a method that evaluates genotype likelihoods to model identity-by-descent, and is specifically designed to account for low coverage data (see Supplementary Text S12, Table S12.1).

Within Szólád we identified four kindreds among the Longobard era burials (gray shadings in Fig 1B), with one particular large one (*Kindred SZ1*, Fig 1B). *Kindred SZ1* (Fig 3A) spans three generations and consists of ten individuals in close spatial proximity. Eight individuals all share recent identity-by-descent (IBD) from SZ24 (one of the oldests individual in the cemetery, between ~45 and 65 years old (yo.)), while another two individuals, SZ15 and SZ19, are more remotely connected genealogically to the kindred via SZ6, a young male aged 8-12 yo. at the time of death. While SZ6 is related to all other individuals in this kindred, sharing the greatest IBD with siblings SZ8 and SZ14, we are unable to determine the exact genealogical relationships involved, likely because of its low SNP coverage (0.048x).

**Fig 3.**
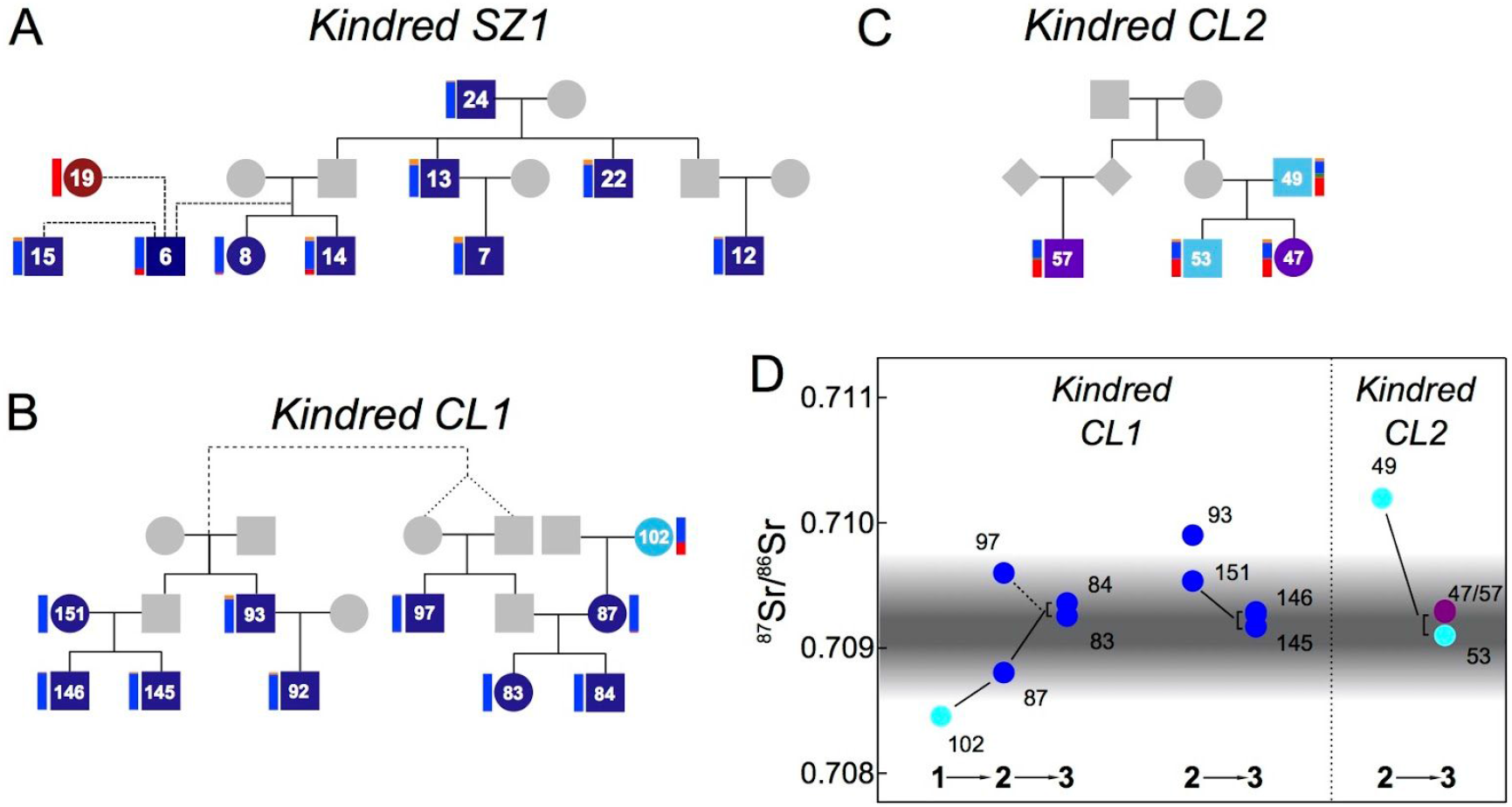
Major kindreds in Szólád and Collegno. *Kindreds SZ1* (A), *CL1* (B) and *CL2* (C), with colors corresponding to criteria and labelling in Fig 2. ADMIXTURE coefficients (vertical bars on side of each individual) estimated using 1000 Genomes20 European populations only. Dashed lines indicate relationships of unknown degree (including past inbreeding in the parents of CL97). Panel D displays strontium isotope ratios for *Kindreds CL1* and *CL2*, where available Estimate of local range at Collegno is shown using black shading, as detailed in Fig. 4. Numbers indicate generations.

Individuals in this kindred were buried with a rich diversity of grave goods, and all but one was buried in elaborate ledge graves. Only two members of this kindred are female, SZ8 and SZ19, who are estimated to be 3-5 and 17-25 yo. respectively at death. The rest are males, aged from ~1 to ~65 yo. These graves occupy a prominent position in the northwest of the cemetery, with all but SZ19 found amongst the core group (she is instead found in the external half-ring of women; see Figure S2.2). Six male individuals in this kindred were buried with weapons, despite three (SZ7, SZ14, SZ15) being teenagers at the time of death (12-17 yo.). The adult males in this kindred (SZ24, SZ13 and SZ22) appear to have had access to a diet particularly high in animal protein, as inferred from nitrogen isotope analysis^13^. Individual SZ13 has the deepest grave and is the only individual in the whole cemetery whose burial includes a weighing scale and a horse, which may be an indication of his differentiated status in that society. We note that this kindred lacks adult female descendants of SZ24, though we were unable to sample some of the female graves in the half-ring structure (unsampled graves SZ21 and SZ29 could still be potential mothers of the third generation; individuals SZ17 and SZ26 can be excluded based on mtDNA analysis by Vai et al. (in review)).

While individuals in this kindred are predominantly of a central/northern European genetic ancestry, they are not genetically homogenous. Again, SZ19 is an outlier, strikingly possessing 100% TSI ancestry with a high probability PAA assignment to the Italian peninsula. In addition SZ6 and the two third generation siblings SZ8 and SZ14 also possess a small but noticeable TSI ancestry component. Assuming that one of the siblings’ parents represented the central European ancestry seen in their two uncles (SZ13 and SZ22), we inferred (using an adapted version of Spatial Ancestry Analysis, SPA, Supplementary Text S13) that the other parent likely possessed an ancestry that most resembled modern day French individuals (Figures S13.5-6). This latter individual would probably have been female, as while SZ14 has a similar Y chromosome to SZ13 and SZ22, both siblings possess a different mtDNA type to their uncles (I3 versus N1b2).

In Collegno, we identified three kindreds, with one particularly extensive one. Nine of the ten individuals from the largest kindred (*Kindred CL1;* Fig 1C and 3B) were buried in elaborate graves and/or with artifacts (Fig 1C). In contrast to Szólád, individuals with close biological relationships occupied spatially distant graves. Interestingly the spatial cluster with six individuals is chronologically older (570/590-610 CE) than the more westerly trio (Figure S3.8). *Kindred CL2* (Fig 3C) is found in the east part of Collegno, with graves positioned in a row running north to south (Fig 1C).

Similar to *Kindred SZ1*, *Kindred CL1* is predominantly of central/northern European ancestry. However, while genetically quite similar, on average members of this kindred possess slightly less FIN ancestry and are thus more shifted towards northwestern Europe in the PCA, SPA analysis and PAA. In addition, this group is again not genetically homogenous, with the unsampled father of CL87 being of much greater central/northern European ancestry than the mother, CL102, who has an ancestry profile again most consistent with modern day France based on PAA (Figures S13.8). *Kindred CL2* also has a more mixed genetic ancestry based on the ADMIXTURE analysis, containing significant contributions from CEU+GBR, TSI and IBS. This would generally associate individuals in this kindred with a more modern central European ancestry than *Kindreds SZ1* and *CL1*. Interestingly the grave goods of the daughter CL47 and an unsampled adjacent female, CL48, resemble burials from this time in southern France and Switzerland (Supplementary Text S3). Members of these two large kindred groups also appear to have generally consumed more animal protein than other individuals in the cemetery, as suggested by nitrogen isotopic analysis (Figures S15.4).

### Genetic ancestry is associated with elements of material culture

In both cemeteries individuals with predominantly central/northern and southern European ancestry possess very distinctive grave furnishings. In order to quantify this relationship, we classified individuals into either Northern (*N*) or Southern (*S*) groups based on their proportion of CEU+GBR+FIN ancestry versus TSI+IBS ancestry (Table S14.1), and used these to conduct a series of Fisher’s exact tests for their association with material culture (we note that our results were robust to our specific ancestry cutoffs; see Supplementary Text S14). We focused on artifacts potentially associated with either specific cultural traditions (e.g. S-brooches and stamped pottery: see details in Supplementary Text S14) or individual profession or status (e.g. war weapons). In both Szólád and Collegno individuals with *N* ancestry were significantly more often buried with grave goods (p-value < 0.0071, Table S14.2). In contrast, no *S* individual was buried with such artifacts, with only two exceptions (females SZ19 and SZ31). This association between genetic ancestry and material culture is particularly significant for beads (from necklaces and pendants) and food offerings in Szólád, as well as weapons in both Collegno and Szólád (Table S14.4). We note that one artifact in grave SZ19 is stylistically distinct (possibly Roman) from the artifacts found in other graves in the same cemetery. Grave type (p-value < 0.02, Table S14.3) also significantly differs between groups in both cemeteries, with *N* individuals presenting graves with wooden elements, as opposed to simple pits (more common amongst graves with *S* individuals).

### Comparison of genetic and strontium isotope data

We also generated new strontium isotope data (^87^Sr/^86^Sr) for Collegno to complement the existing data from Szólád^13^ and analyzed them within the context of our genomic data in order to better understand patterns of immigration to these two sites (Supplementary Text S15, Table S15.1). Within Szólád we find that adult individuals with both predominantly central/northern and southern genomic ancestry possess similar non-local signatures (Alt et al.^13^ described this as Range I) (Fig 4). This might suggest that individuals from both ancestry groups immigrated into Szólád together despite the differences in material culture. However, we also note generally a quite diverse non-local range amongst adults with central/northern ancestry (for example SZ4 and SZ16 are extreme outliers), pointing to not all individuals having origins from the same location prior to settling in Szólád.

**Fig 4.**
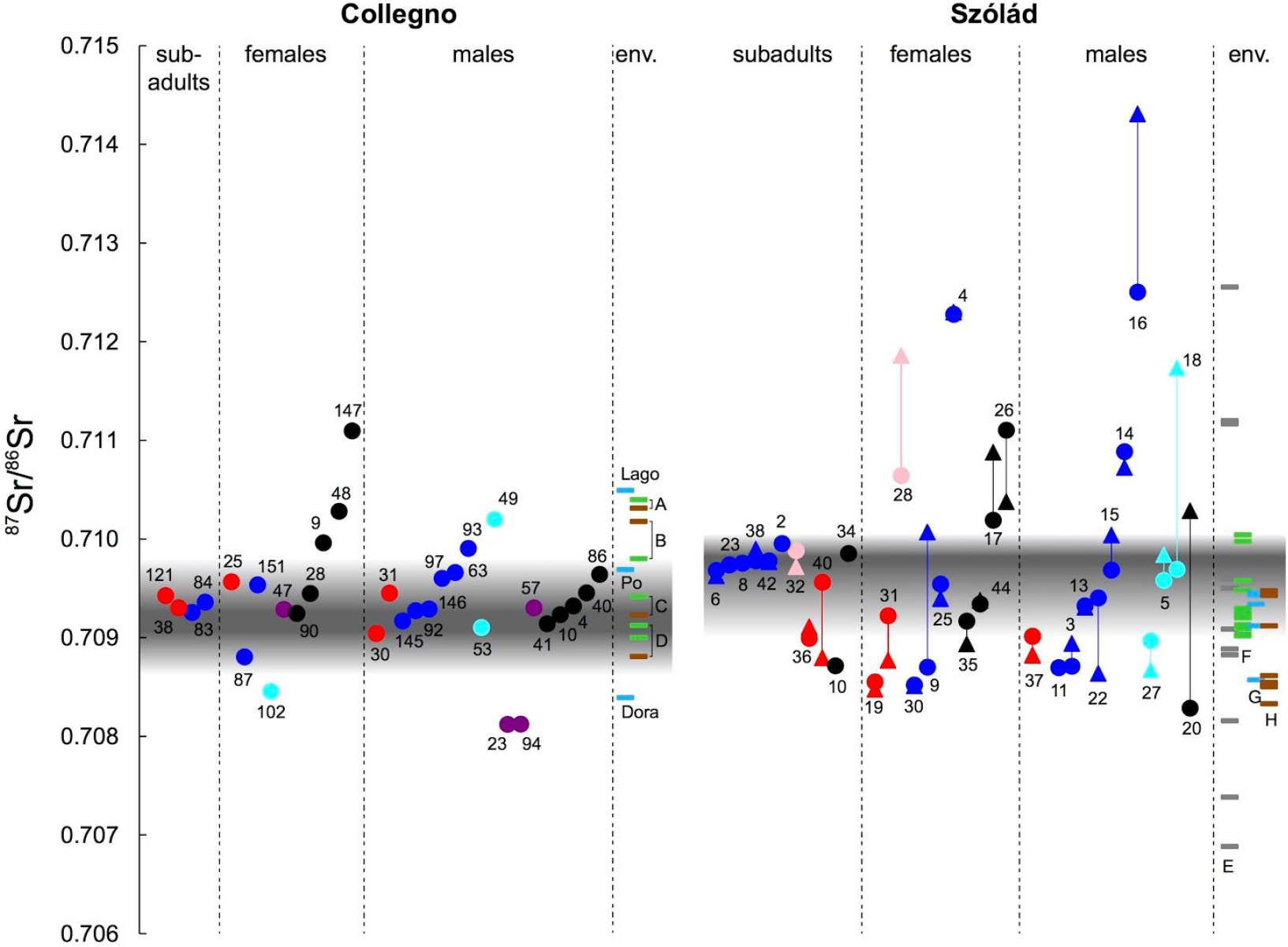
Strontium isotope ratios in Szólád and Collegno. Evidence for migrants at Collegno and Szólád, as suggested by 87Sr/86Sr values of human tooth enamel and environmental reference samples. The grey band denotes the local bioavailable strontium isotope range, with individuals falling outside of that considered non-local. A: Collino di Superga (clays and marls); B: Lago Piccolo di Avigliana (glacial deposits); C: Castello di Avigliana (serpentinite, green schist); D: Collegno site (Pleistocene gravels); E: Bakony Mountains; F: vegetation south of Lake Balaton; G: water south of Lake Balaton; H: soil near Szólád (Szólád data from 13). Colors corresponding to criteria and labelling in Fig 2, with black samples being those individuals lacking genomic data.

In contrast, in Collegno it was notable that the five individuals with major southern ancestry are primarily assigned to Italy using PAA, exhibited local strontium signatures. When examining the two major kindred, we observe the striking general pattern that earlier generations had strontium isotope values that diverged from the local range more than later generations (Fig 3D, Figure S15.3). This appears to fit a model of individuals of central/northern European ancestry migrating and settling in Collegno amongst a set of local individuals of primarily Italian origin.

## DISCUSSION

The most striking feature of our data is the inference of two main clusters of genetic ancestry that are shared amongst our two 6th-7th century cemeteries separated by almost 1,000 km. If we orientate this ancestry using modern populations they correspond to an axis of central/northern European ancestry versus southern European ancestry. In both Szólád and Collegno this genetic variation mirrors the variation that emerges from their mortuary practices, i.e. how living members of the community represented the individuals that they buried. This perhaps suggests that the concept of long-term shared common descent in shaping social identity, as reflected in the material culture, may have had an actual biological basis in these two cemeteries. However, whether the association between genetic ancestry and material culture reflects specific peoples mentioned in historical texts (i.e. Longobards) or stemmed from a deeper/long-term descent (of mixed barbarian ancestries) is as yet unclear.

**Social Organization During Migration versus Settlement.** Our genomic characterization of Szólád and Collegno provides novel insight into the structures and hierarchies of societies from the Migration Period in two very different contexts. On the one hand a previous study of Szólád had identified this cemetery as belonging to a highly mobile community that was settled in the region for approximately only one generation. We were able to show in particular that the burials in this cemetery were organized predominantly around a three-generation kindred with ten members (*Kindred SZ1*), all but one of which were of predominantly central/northern ancestry. Members of this kindred stand out in relation to others for a number of reasons: (i) access to diet higher in animal protein; (ii) graves occupying a prominent position in the cemetery, (iii) the presence of the oldest individual in the cemetery (SZ24), and (iv) the individual (SZ13) with the deepest grave and the only one buried with a horse, Thuringian type pottery, and a scale with weights and coins.

Surrounding this kindred within the area demarcated by the half-ring of women there are 10 males. Of the nine for which we have genetic data, all have predominantly central/northern European genetic ancestry but some have both additional TSI and IBS ancestry (for example SZ5), suggesting somewhat diverse genetic and thus possibly different geographic origins amongst each other and from the focal *Kindred SZ1*. All adults and teenagers have weapons and three out of four adults share the same non-local Sr signature as adults in *Kindred SZ1*. The half-ring of women itself has a mixture of individuals (10 adults and 1 child), with five that have predominantly central/northern ancestry, three that have majority TSI+IBS ancestry and they have a wide range of strontium isotope ratios. SZ19 is found amongst this half-ring and though she is part of *Kindred SZ1*, she lacks the ledge graves of the other members, has a distinct material culture and is of southern ancestry. To what extent her lack of IBD with SZ24 and her different ancestry contributed to her more peripheral burial compared to both the female SZ8 (who is buried within the core) and the male SZ15 (who similarly unrelated to SZ24 but again found within the core) is an open question. Analysis of more cemeteries from the period will be required to better understand such complexities.

Since the adults were almost all non-local, it is tempting to suggest that we may be observing the historically described *fara* during migration. Regardless, this group appears to be a unit organized around one high-status, kin-based group of predominantly males, but also incorporating other males that may have some common central/northern European descent. The relative lack of adult female representatives from *Kindred SZ1*, the diverse genetic and isotope signatures of the sampled women around the males and their rich graves goods suggests that they may have been acquired and incorporated into the unit during the process of migration (perhaps hinting at a patrilocal societal structure that has been shown to be prominent in Europe during earlier periods^31^).

The remaining part of this community for which we have genomic data (N=7) is composed of individuals of mainly southern European genetic ancestry that are conspicuously lacking grave goods and occupy the southeastern part of the cemetery, with randomly oriented graves with straight walls. While the lack of grave goods does not necessarily imply that these individuals were of lower status, it does point to them belonging to a different social group. Interestingly, the strontium isotope data suggest that they may have migrated together with the warrior-based group from outside Szólád, but barriers to gene flow were largely been maintained.

In contrast to Szólád, Collegno likely reflects a community that eventually settled for multiple generations. Organization around at least one large extended immigrant kindred once again seems to have been a key element of social organization. However, there is more spatial variation, with the kindred spreading outwards from the center point of the cemetery over time. There is also one other significant but smaller immigrant kindred of a different genetic origin and material culture that also holds a central position in the cemetery, and, for this first period of occupation at least, these two groups appear to remain distinct genetically despite living in the same location. Unlike at Szólád, we find evidence of only one other individual (CL63) that is of predominant central/northern ancestry that does not belong to the major kindred unit (though our sampling of this cemetery is not as complete as Szólád). On the other hand, individuals of southern ancestry of the type that would typically be found in the region today appear to be local to the Collegno area based on isotope data, show much more scattered burials and are poorer with regard to grave goods and animal protein consumption. As such it is tempting to infer a scenario of these large immigrant barbarian families exerting a dominating influence on the original resident population.

### Evidence for Migrating Barbarians and “Longobards”

Our two cemeteries overlap chronologically with the historically documented migration of Longobards from Pannonia to Italy at the end of the 6th century. It is thus intriguing that we observe that central/northern European ancestry is dominant not only in Szólád, but also in Collegno. Based on modern genetic data we would not expect to see a preponderance of such ancestry in either Hungary or especially Northern Italy. While we do not yet know the general genomic background of Europe in these geographic regions just before the establishment of Szólád and Collegno, other Migration Period genomes from the UK and Germany show a fairly strong correlation with modern geography (while also possessing a similar central/northern European ancestry component to that found in Szólád and Collegno). Going further back in time, Late Bronze Age Hungarians show almost no resemblance to populations from modern central/northern Europe, especially compared to Bronze Age Germans and in particular Scandinavians, who, in contrast, show considerable overlap with our Szólád and Collegno central/northern ancestry samples. Coupled with the strontium isotope data, our paleogenomic analysis suggest that the earliest individuals of central/northern ancestry in Collegno were probably migrants while those with southern ancestry were local residents. Our results are thus consistent with an origin of barbarian groups such as the Longobards somewhere in Northern and Central Europe east of the Rhine and north of the Danube. Thus our results cannot reject the migration, its route, and settlement of “the Longobards” described in historical texts.

We note however that whether these people identified as “Longobard” or any other particular barbarian people is impossible to assess. Modern European genetic variation is generally highly structured by geography^22,32^, even at the level of individual villages^33^. It is, therefore, surprising to find significant diversity, even amongst individuals with central/northern ancestry, within small, individual Langobard cemeteries. Even amongst the two family groups of primarily central/northern ancestry, who may have formed the heart of such migration, there is clear evidence of admixture with individuals with more southern ancestry. If we are seeing evidence of movements of barbarians, there is no evidence that these were genetically homogenous groups of people.

### Conclusions

We have observed an intriguing association between genetic evidence, isotope data and material culture that sheds new light on the social organization of sixth century communities during both migration and settlement phases. Our study reveals for the first time the importance of biological relatedness in shaping social relationships. It also potentially supports long-distance barbarian migrations described in historical texts. A key aspect of our approach is the in-depth sampling of entire cemeteries. We propose that this is a conceptual and methodological advance: In the future one must use a similar whole cemetery-based genomic methodology to explore whether the results observed here are common to other Longobard-era cemeteries and to other sites from Late Antiquity and the Early Middle Ages. The genetic complexity observed within these cemeteries presents a new set of questions concerning population structure within past societies. Moreover, if the genetic similarity we observe between Szólád and Collegno appears in other contemporary cemeteries, we will be able to appreciate with greater confidence the extent and dynamics of these movements and of the invasion of barbarian peoples across the Roman Empire.

## METHODS

### DNA isolation, screening, sequencing and bioinformatic processing

Bone specimens from Szólád and Collegno were prepared in clean-room facilities dedicated to ancient DNA in the Laboratory of Molecular Anthropology and Paleogenetics, University of Florence. DNA extraction was performed using a silica-based protocol^34^. Genomic libraries were prepared at the University of Florence and at the Max Planck Institute for the Science of Human History in Jena according to modified Illumina protocols and were screened for endogenous human DNA via mapping to the human reference genome after re-sequencing. DNA postmortem damage (PMD) patterns typical of ancient DNA were assessed with MapDamage^35^. The 10 samples (all from Szólád) with best DNA quality and concentration were submitted for WGS on an Illumina HiSeq 2500 1TB. Another 53 samples underwent a capture and NextSeq sequencing for ~1.2M SNPs for a targeted average coverage of ~1x per sample. Reads were trimmed, merged (where applicable), mapped and filtered for PCR duplications according to a protocols optimized for ancient DNA^36,37^. Genotype likelihood estimation and pseudo-haploid and diploid calling was performed using in-house Python scripts (www.github.com/kveeramah). For samples subject to partial UDG treatment, genotype likelihood estimation was implemented, ignoring the first and last three bases of reads. Genotype calling considered PMD for the remaining samples^38^. Finally, a pseudo-haploid call set was generated by sampling a random read at each position. See Supplementary Text S4 and S5 for further details.

### Modern and Ancient Reference Samples

We assembled SNP data matching the 1240K capture for three modern reference datasets and one ancient reference dataset for comparison to the early Medieval samples generated in this study. The three modern reference sets consisted of an imputed^39^ European POPRES^40^ SNP set with ~300K SNPs, an imputed Eurasian23 SNP set with ~700K, and a 1000 Genomes^20^ and SGDP^21^ whole genome set matching all 1240K SNPs. Our ancient reference set consisted of 211 ancient West Eurasians from 6,300 − 300 BC^19^ with calls made at the 1240K capture SNPs. Depending on the context, we made pseudo-haploid calls by randomly drawing one allele from a diploid genotype.

### Biological Relatedness Inference

We used the software lcMLkin^30^ to estimate biological relatedness and coancestry coefficients for every pair of samples. In addition to using the allele frequencies of the ancient samples themselves, we also adapted the software to utilize allele frequencies from other sources, in this case the CEU and TSI 1000 Genomes populations, and incorporate admixture.

### Principal Component Analysis

PCA of SNP data was conducted using smartpca^41^. When analyzing Migration Period individuals against reference populations, individual pseudo-haploid PCAs were conducted for each ancient sample separately, and individual analyses were then combined using a Procrustes transformation in `R` using the `vegan` package as described previously^42^. LD pruning was performed using the --indep-pairwise function in PLINK^43^.

### Model-based Clustering Analysis

Supervised model-based clustering was performed using ADMIXTURE^26^. Dependent on the analysis, target Migration Period samples were analyzed individually (to avoid the effects of relatedness) or together (using a set of unrelated individuals that maximized SNP number)

### Population Assignment Analysis

PAA was conducted as described in Veeramah et al.^25^ using the POPRES and HellBus datasets.

### Spatial Ancestry Analysis

We applied the software SPA^44^ to analyse the POPRES imputed SNP dataset. We also further extended the software to allow the use of pseudo-haploid calls in our ancient samples, and to infer the location of one parent of an admixed individual given the known location of another parent.

### Rare Variant Analysis

We followed the approach of Schiffels et al.^24^ to examine the relative sharing of central/northern European and southern European-specific rare variants, and used the software `rarecoal` to assign our ancient whole genomes to a branch on a bifurcating demographic model based on the analysis of rare variants in modern European populations (TSI, IBS and GBR from the 100 Genomes project^20^, Denmark^45^ and the Netherlands^46^.

### Y chromosome Analysis

The phylogenetic position of each Y chromosome variant observed in the whole sample was established according to its occurrence in public database or in the published literature^47,48^. The lack of base calls due to the absence of reads at a position in a particular sample was resolved either as an ancestral or derived allele by a hierarchical inferential method according to the phylogenetic context based on a cladistic approach. The phylogenetically informative SNPs were used to build a parsimony-based phylogenetic tree using the Pars application of the `Phylip v3.69` package. `FigTree v1.4.2` software was used to display the generated tree.

### Isotope Analysis

Strontium, oxygen, carbon and nitrogen isotope analysis was carried out on 33 first-period individuals from Collegno, together with faunal and environmental reference samples. Tooth enamel powder primarily from second premolars or molars was used for strontium isotope analysis at the Isotope Geochemistry Laboratory of the Department of Earth Sciences, University of Cambridge. Preparation of tooth enamel powder for oxygen isotope analysis followed the method described by Balasse et al.^49^. Collagen was extracted from bones for nitrogen and carbon isotope analysis, based on the method detailed by Privat et al^50^, These analyses were carried out at the Godwin Laboratory, University of Cambridge. To determine local bioavailable strontium values, samples of water, vegetation and soil were collected both at the site of the cemetery of Collegno and at locations considered to be about a day’s walk from the site and then pre-treated following the procedure outlined in Maurer et al.^51^. Faunal samples were taken from six species from a 3rd/4th century site in Piazza Castello in Turin to provide an ecological baseline for human diet.

### Data availability

Sequencing data (as processed BAM files) are available from the NCBI sequence read archive (SRA) database under accession # SRP132561 (1240 capture data) and SRP132581 (WGS data). Code generated to call variants in the ancient samples is available at: https://github.com/kveeramah/

## ACKNOWLEDGEMENTS

We thank Kurt Alt for his role in scientifically characterizing Szólád, and Kathryn Twiss for her comments on the manuscript. We are particularly appreciative of discussions with Philipp Von Rummel, Falko Daim, Frans Theuws and Helmut Reimitz. We are grateful to Hazel Chapman and James Rolfe for help with isotopic analyses and to David Redhouse for help with an illustration. We thank Christoph Lippert, Amalio Talenti, Haibao Tang, and Emily Wong for supplementary analyses not included in this version of the manuscript. We thank Marta Burri and lab members of the Max Planck Institute for the Science of Human History in Jena for laboratory support. This work was supported by National Science Foundation award #1450606, the Anneliese Maier Research Award of the Alexander von Humboldt Foundation, the Max Planck Society, the German Federal Ministry for Education and Research, the Swedish Riksbankens Jubieleumfond, and the Gerard B. Lambert Foundation, the Italian Ministry for University and Research Department of Excellence Program.

## AUTHOR CONTRIBUTIONS

I.K, L.P.B, E.B, C.G, T.V, D.W and U.F. provided the archaeological material and/or performed the archaeological analysis and interpretation. M.C.LR, W.P and P.J.G provided the historical background and interpretation. S.V, C.P, A.M, B.M, M.L, J.K and D.C. performed the ancient DNA lab work and screening. C.E.G.A, D.B, C.P, S.G, G.B and K.R.V and performed the downstream bioinformatics and population genetic analysis. P.A. performed the Y-chromosome analysis. S.H. performed the isotope analysis. C.E.G.A, P.J.G and K.R.V wrote the paper. W.P, P.J.G and K.R.V conceived the study.

## REFERENCES

1. Pohl, W. & Reimitz, H. Strategies of distinction: the construction of ethnic communities, 300–800. 2, (Brill, 1998).

2. Geary, P. The Myth of Nations: The Medieval Origins of Europe. (Princeton University Press, 2003).

3. Heather, P. The Fall of the Roman Empire: A New History of Rome and the Barbarians. (Oxford University Press, 2005).

4. Werner, J. Zur Entstehung der Reihengräberzivilisation. in Siedlung, Sprache und Bevölkerungsstruktur im Frenkenreich (ed. Petri, F.) 285–325 (Darmstadt: Wissenschaftliche Buchgesellschaft, 1973).

5. Bierbrauer, V. Archäologie der Langobarden in Italien: ethnische Interpretation und Stand der Forschung. in Die Langobarden. Herrschaft und Identität (eds. Pohl, W. & Erhart, P.) Forschungen zur Geschichte des Mittelalters 9, 21–66 (Wien: Verlag der Österreichischen Akademie der Wissenschaften., 2005).

6. Brather, S. Ethnische Interpretationen in der frühgeschichtlichen Archäologie. Geschichte, Grundlagen und Alternativen. 78, (Ergänzungsband 42, 2004).

7. Ward-Perkins, B. The fall of Rome and the end of civilization. (Oxford University Press, Oxford, 2005).

8. Halsall, G. Barbarian migrations and the Roman West, 376–568. (Cambridge University Press, Cambridge, 2007).

9. Jarnut, J. Geschichte der Langobarden. (Kohlhammer, 1982).

10. Ausenda, G., Delogu, P. & Wickham, P. The Langobards before the Frankish Conquest: An Ethnographic Perspective. (Boydell Press, 2009).

11. Pohl, W. & Erhart, P. Die Langobarden Herrschaft und Identität (Denkschriften der philosophisch-historischen Klasse). 9, (Verlag der österreichischen Akademie der Wissenschaften, 2005).

12. Murray, A. C. Germanic Kinship Structure: Studies in Law and Society in Antiquity and in the Early Middle Ages. (Pontifical Institute of Mediaeval Studies, Toronto, 1983).

13. Alt, K. W. et al. Lombards on the move--an integrative study of the migration period cemetery at Szólád, Hungary. PLoS One 9, e110793 (2014).

14. Koncz, I. 568 — A historical date and its archaeological consequences. Acta Archaeologica Academiae Scientiarum Hungaricae 66, 315–340 (2015).

15. Pejrani Baricco, L. Presenze longobarde: Collegno nell’alto medioevo; [Collegno, Certosa Reale, 18 aprile - 20 giugno 2004]. (2004).

16. Pinhasi, R. et al. Optimal Ancient DNA Yields from the Inner Ear Part of the Human Petrous Bone. PLoS One 10, e0129102 (2015).

17. Rohland, N., Harney, E., Mallick, S., Nordenfelt, S. & Reich, D. Partial uracil-DNA-glycosylase treatment for screening of ancient DNA. Philos. Trans. R. Soc. Lond. B Biol. Sci. 370, 20130624 (2015).

18. Fu, Q. et al. An early modern human from Romania with a recent Neanderthal ancestor. Nature 524, 216–219 (2015).

19. Mathieson, I. et al. Genome-wide patterns of selection in 230 ancient Eurasians. Nature 528, 499–503 (2015).

20. 1000 Genomes Project Consortium et al. A global reference for human genetic variation. Nature 526, 68–74 (2015).

21. Mallick, S. et al. The Simons Genome Diversity Project: 300 genomes from 142 diverse populations. Nature 538, 201–206 (2016).

22. Novembre, J. et al. Genes mirror geography within Europe. Nature 456, 98–101 (2008).

23. Hellenthal, G. et al. A genetic atlas of human admixture history. Science 343, 747–751 (2014).

24. Schiffels, S. et al. Iron Age and Anglo-Saxon genomes from East England reveal British migration history. Nat. Commun. 7, 10408 (2016).

25. Veeramah, K. R. et al. Population genomic analysis of elongated skulls reveals extensive female-biased immigration in Early Medieval Bavaria. Proc. Natl. Acad. Sci. U. S. A. (in press).

26. Alexander, D. H., Novembre, J. & Lange, K. Fast model-based estimation of ancestry in unrelated individuals. Genome Res. 19, 1655–1664 (2009).

27. Meyer, C., Ganslmeier, R., Dresely, V. & Alt, K. W. New approaches to the reconstruction of kinship and social structure based on bioarchaeological analysis of Neolithic multiple and collective graves. in Theoretical and Methodological Considerations in Central European Neolithic Archaeology (eds. Kolář, J. & Trampota, F.) 2325, 11–23 (2012).

28. Haak, W. et al. Ancient DNA, Strontium isotopes, and osteological analyses shed light on social and kinship organization of the Later Stone Age. Proc. Natl. Acad. Sci. U. S. A. 105, 18226–18231 (2008).

29. Johnson, K. M. & Paul, K. S. Bioarchaeology and Kinship: Integrating Theory, Social Relatedness, and Biology in Ancient Family Research. J Archaeol Res 24, 75–123 (2016).

30. Lipatov, M., Sanjeev, K., Patro, R. & Veeramah, K. Maximum likelihood estimation of biological relatedness from low coverage sequencing data. bioRxiv. (2015).

31. Knipper, C. et al. Female exogamy and gene pool diversification at the transition from the Final Neolithic to the Early Bronze Age in central Europe. Proc. Natl. Acad. Sci. U. S. A. 114, 10083–10088 (2017).

32. Leslie, S. et al. The fine-scale genetic structure of the British population. Nature 519, 309–314 (2015).

33. O’Dushlaine, C. et al. Genes predict village of origin in rural Europe. Eur. J. Hum. Genet. 18, 1269–1270 (2010).

34. Dabney, J. et al. Complete mitochondrial genome sequence of a Middle Pleistocene cave bear reconstructed from ultrashort DNA fragments. Proc. Natl. Acad. Sci. U. S. A. 110, 15758–15763 (2013).

35. Ginolhac, A., Rasmussen, M., Gilbert, M. T. P., Willerslev, E. & Orlando, L. mapDamage: testing for damage patterns in ancient DNA sequences. Bioinformatics 27, 2153–2155 (2011).

36. Kircher, M. Analysis of high-throughput ancient DNA sequencing data. Methods Mol. Biol. 840, 197–228 (2012).

37. Peltzer, A. et al. EAGER: efficient ancient genome reconstruction. Genome Biol. 17, 60 (2016).

38. Botigué, L. R. et al. Ancient European dog genomes reveal continuity since the Early Neolithic. Nat. Commun. 8, 16082 (2017).

39. Das, S. et al. Next-generation genotype imputation service and methods. Nat. Genet. 48, 1284–1287 (2016).

40. Nelson, M. R. et al. The Population Reference Sample, POPRES: a resource for population, disease, and pharmacological genetics research. Am. J. Hum. Genet. 83, 347–358 (2008).

41. Patterson, N., Price, A. L. & Reich, D. Population structure and eigenanalysis. PLoS Genet. 2, e190 (2006).

42. Veeramah, K. R. & Hammer, M. F. The impact of whole-genome sequencing on the reconstruction of human population history. Nat. Rev. Genet. 15, 149–162 (2014).

43. Purcell, S. et al. PLINK: a tool set for whole-genome association and population-based linkage analyses. Am. J. Hum. Genet. 81, 559–575 (2007).

44. Yang, W.-Y., Novembre, J., Eskin, E. & Halperin, E. A model-based approach for analysis of spatial structure in genetic data. Nat. Genet. 44, 725–731 (2012).

45. Besenbacher, S. et al. Novel variation and de novo mutation rates in population-wide de novo assembled Danish trios. Nat. Commun. 6, 5969 (2015).

46. Genome of the Netherlands Consortium. Whole-genome sequence variation, population structure and demographic history of the Dutch population. Nat. Genet. 46, 818–825 (2014).

47. Francalacci, P. et al. Low-pass DNA sequencing of 1200 Sardinians reconstructs European Y-chromosome phylogeny. Science 341, 565–569 (2013).

48. Hallast, P. et al. The Y-chromosome tree bursts into leaf: 13,000 high-confidence SNPs covering the majority of known clades. Mol. Biol. Evol. 32, 661–673 (2015).

49. Balasse, M. & Tresset, A. Early Weaning of Neolithic Domestic Cattle (Bercy, France) Revealed by Intra-tooth Variation in Nitrogen Isotope Ratios. J. Archaeol. Sci. 29, 853–859 (2002).

50. Privat, K. L., O’connell, T. C. & Richards, M. P. Stable Isotope Analysis of Human and Faunal Remains from the Anglo-Saxon Cemetery at Berinsfield, Oxfordshire: Dietary and Social Implications. J. Archaeol. Sci. 29, 779–790 (2002).

51. Maurer, A.-F. et al. Bioavailable 87Sr/86Sr in different environmental samples--effects of anthropogenic contamination and implications for isoscapes in past migration studies. Sci. Total Environ. 433, 216–229 (2012).

